# Effects of two seminal fluid proteins on post-mating behavior in the simultaneously hermaphroditic flatworm *Macrostomum lignano*

**DOI:** 10.1101/734301

**Authors:** Michael Weber, Bahar Patlar, Steven A. Ramm

## Abstract

Along with sperm, in many taxa male ejaculates also contain a complex mixture of proteins, peptides and other substances found in seminal fluid. Once seminal fluid proteins (SFPs) are transferred to the mating partner, they play crucial roles in mediating post-mating sexual selection, since they can modulate the partner’s behavior and physiology in ways that influence the reproductive success of both partners. One way in which sperm donors can maximize their own reproductive success is by changing the partners’ (sperm recipient’s) postcopulatory behavior to prevent or delay re-mating, thereby decreasing the likelihood or intensity of sperm competition. We therefore adopted a quantitative genetic approach combining gene expression and behavioral data to identify candidates that could mediate such a response in the simultaneously hermaphroditic flatworm *Macrostomum lignano*. We identified two putative SFPs - Mlig-pro46 and Mlig-pro63 - that exhibit a negative genetic correlation between transcript expression and mating frequency. Importantly, however, in one of the two different group sizes, differing in their sperm competition level, in which we measured genetic correlations, these same two transcripts are also linked to a second post-mating behavior in *M. lignano*, namely the ‘suck’ behavior of recipients in which, upon ejaculate receipt, the worm places its pharynx over its female genital opening and appears to attempt to remove ejaculate components. To therefore investigate directly whether these two candidates manipulate partner behavior, and test whether this impacts on competitive fertilization success, we performed a manipulative experiment using RNA interference-induced knockdown to ask how loss of Mlig-pro46 and Mlig-pro63 expression, singly and in combination, affects mating frequency, partner suck propensity and both defensive and offensive sperm competitive ability (*P*_1_ and *P*_2_, respectively). None of the knock-down treatments impacted strongly on mating frequency or sperm competitive ability, but the knock-down of Mlig-pro63 resulted in a significantly decreased ‘suck’ propensity of mating partners. This suggests that Mlig-pro63 may normally act as a cue in the ejaculate to trigger recipient suck behavior, though the functional and adaptive significance of these two seminal proteins from a donor perspective remains enigmatic.

## Introduction

In polyandrous species, when females store sperm from multiple males, post-mating sexual selection can occur if sperm from different males compete with each other for fertilization (i.e. sperm competition; (Parker 1970)) and/or females choose specific sperm to fertilize eggs (i.e. cryptic female choice; (Thornhill 1983; Eberhard 1996)). As a consequence, several male adaptations have evolved to increase their fertilization success, including displacing sperm of previous males (Waage 1986; Harshman and Prout 1994; Marie-Orleach et al. 2014), preventing females from re-mating (Radhakrishnan and Taylor 2007, 2008; Uhl et al. 2009; Abraham et al. 2016) or possessing morphologically more competitive sperm (Birkhead 1995; Birkhead and Pizzari 2002). This can lead to a sexual conflict over the optimal fitness strategies concerning reproduction, often resulting in cycles of sexually antagonistic coevolution (Parker 1979; Chapman et al. 2003; Arnqvist and Rowe 2005; Chapman 2018).

The seminal fluid proteins (SFPs) found in the ejaculate play crucial roles in reproduction and can modulate the mating partner’s behavior and physiology such that they affect the reproductive success of both partners (reviewed in: Chapman 2001; Avila et al. 2011; Sirot et al. 2015; Hopkins et al. 2017), making these proteins key mediators of post-mating sexual selection (Chapman 2001; Hodgson and Hosken 2006; Poiani 2006; Cameron et al. 2007; Ram and Wolfner 2007a). And – because they can modulate reproductive behavior and physiology in many ways to the advantage of the seminal fluid-donating individual, but not necessarily that of the seminal fluid-receiving individual – sexual conflict (Chapman et al. 1995b; Arnqvist and Rowe 2005; Sirot et al. 2015).

SFPs influence subsequent female physiology and behavior in various ways (reviewed in: Poiani 2006; Avila et al. 2011; Sirot et al. 2015; Hopkins et al. 2017). For example, specific SFPs are known to modulate egg production, ovulation, and/or egg-laying rates (Gillott 2003; Poiani 2006; Ram and Wolfner 2007a; Avila et al. 2011). Other SFPs can affect sperm storage (Neubaum and Wolfner 1999; Chapman et al. 2000; Qazi 2003).

One example of a potential sexual conflict mediated through seminal fluid is a mating-induced change of the partners’ sexual receptivity. Decreased sexual receptivity of mated females occurs in a wide range of insects, and it is suggested that males benefit from this change because it decreases the likelihood or intensity of sperm competition. In *D. melanogaster*, mated females actively reject courting males and the SFP sex peptide received with the ejaculate plays a central role in inducing this change (Chapman et al. 2003; Liu and Kubli 2003; Yapici et al. 2008; Häsemeyer et al. 2009; Ram and Wolfner 2009; Yang et al. 2009): females mated to sex peptide null males remain highly receptive to re-mating (Chapman et al. 2003; Liu and Kubli 2003).

Although *Drosophila* has been the main model species to date for studying such seminal fluid-mediated effects, there are several other reported cases in a broad range of insects where females show a similarly reduced sexual receptivity after receiving full ejaculates or specific SFPs (reviewed in: Simmons 2001; Avila et al. 2011). For example, females of the queensland fruit fly *Bactrocera tryoni* show no difference in sexual receptivity when mated to irradiated males compared with mates of nonirradiated males, suggesting that components of the seminal fluid, not sperm, are responsible for the decreased post-mating receptivity observed (Harmer et al. 2006). Moreover, the injection of male reproductive tract extracts into *B. tryoni* females leads to a decrease in sexual receptivity and shorter copulation times when subsequently mated, similar to behaviors seen in previously mated females (Radhakrishnan and Taylor 2007). In *Anopheles gambiae*, injection of male accessory gland homogenates into virgin females results in a decreased likelihood of re-mating (Shutt et al. 2010). There is also growing evidence that SFPs have important effects on female post-mating behavior in lady beetles (Perry and Rowe 2008a,b), seed beetles (Moya-Laraño and Fox 2006; Rönn et al. 2006; Yamane et al. 2008) and ground beetles (Takami et al. 2008). There is also some evidence for similar seminal-fluid mediated effect in vertebrates, because for example the beta-endorphin found in rat seminal fluid has the capacity to suppress female receptivity (Forsberg et al. 1990). Depending on any direct or indirect benefits females might accrue from additional matings, such effects may often be counter to female interests (Chapman et al. 1995a; Wigby and Chapman 2005; Hollis et al. 2019).

Charnov (1979) first recognized that sexual conflict also occurs in hermaphrodites, since sperm donors can also potentially benefit from transferring ejaculate components manipulating sperm recipients and affecting the outcome of sperm competition (Charnov 1979; Michiels 1998; Koene 2006; Schärer et al. 2015). Moreover, as we explain in more detail below, simultaneous hermaphroditism might also create unique targets for seminal fluid action (Charnov 1979; Schärer et al. 2015; Schärer and Ramm 2016). Such seminal fluid-mediated effects, with a potential beneficial effect for one but a potential harmful effect for the other partner, have already been detected in the pond snail *Lymnaea stagnalis*. The intravaginal injection of one *L. stagnalis* SFP (LyAcp10) led to a decrease in egg laying (Koene et al. 2010) and the injection of two other SFPs (LyAcp8b and LyAcp5) reduced the number of sperm transferred by the recipient in a subsequent mating as a donor, and as a result, decreased their paternity success (Nakadera et al. 2014; see also Schärer 2014). This latter effect emphasizes that it is a potentially adaptive strategy in simultaneous hermaphrodites to steer your partner away from its male function, and that the action of seminal fluid may be one means of doing so (Schärer and Ramm 2016). There is also evidence for the manipulation of the re-mating frequency in a simultaneously hermaphroditic species, namely the love-dart shooting snail *Euhadra quaesita*. Before exchanging sperm, both mating partners attempt to drive their mucous-coated love-dart into their respective partner. In stabbed snails, the intermating interval is longer than in not-stabbed individuals, and so snail pairs injected with mucous subsequently mate less often than control pairs (Kimura et al. 2013).

In our model species *Macrostomum lignano*, the complement of seminal fluid proteins has only just been characterized (Weber et al. 2018). Nevertheless, there are already some indications for potential effects of SFPs. An initial screen of 18 putative SFPs selected without any prior functional information revealed that the RNAi-induced knockdown of at least some SFPs may modulate aspects of fertility and sperm competitiveness (Weber et al, submitted). However, given that this screen identified only a minority of candidate SFPs modulating fertility or sperm competitiveness, and that all of these effects have not yet been verified after controlling for multiple testing, this suggests that more targeted methods may be beneficial to first prioritise candidates mediating specific effects. Patlar et al. (submitted) adopted precisely such an approach, using quantitative genetics to identify six candidate SFPs likely to affect the post-mating ‘suck’ behavior of sperm recipients, based on a negative genetic correlation between SFP transcript expression (Patlar et al. 2018) and suck propensity. The suck behavior in *M. lignano* is a striking response to ejaculate recipient, in which the worm places its pharynx over its female genital opening and appears to attempt to suck out its contents, suggesting this trait has evolved in the context of sexual conflict over control of the received ejaculate (Schärer et al. 2004a; Vizoso et al. 2010; Scharer et al. 2011; Marie-Orleach et al. 2013). RNAi-induced knockdown of two of the six SFP candidate genes, Mlig-pro31 and Mlig-pro32, caused a substantially increased suck propensity of mating partners, suggesting that these proteins indeed manipulate mating partners and may thereby mediate sexual conflict over ejaculate fate, but no clear effect on paternity outcomes in a standardized defensive sperm competition assay (Patlar et al., submitted).

The quantitative genetic approach adopted by Patlar et al. (submitted) provides a powerful framework to identify candidates for subsequent screening of SFP-mediated effects in a targeted manner. We therefore here adopted this same method, enabling us to identify two candidate SFPs, Mlig-pro46 and Mlig-pro63, potentially affecting two behavioural phenotypes of interest: partner re-mating behavior and the propensity for partners to exhibit the suck behavior (see Methods). In order to test whether these proteins in seminal fluid indeed influence these behaviours and have downstream impacts on fitness, we then used RNAi to test these two putative SFPs transcripts for their effects on mating frequency in pairings with untreated partners, on the suck propensity of mating partners, as well as sperm competitive ability. Both putative SFPs were already included in the above-mentioned naïve RNAi screen (Weber et al., submitted), in which Mlig-pro46 showed some evidence for a reduced offensive sperm competitive ability (*P*_2_, see below), making it an even more promising candidate for more detailed characterization.

## Methods

### Study organism and experimental subjects

*M. lignano* is a free-living, outcrossing simultaneous hermaphrodite found in the Northern Adriatic Sea and Eastern Mediterranean (Ladurner et al. 2005; Zadesenets et al. 2016). The worms reach ca. 1.5 mm in body length and the testes and ovaries are located in the central part of the body on either side of a medial gut. The male copulatory organ, the female reproductive organ and the prostate gland cells (where seminal fluid is produced), are all located in the posterior part of the worms (Hyman 1951; Weber et al. 2018). *M. lignano* is obligatory outcrossing, i.e., self-fertilization does not occur, and copulations are reciprocal: both partners receive and donate sperm at the same time (Schärer and Ladurner 2003). *Macrostomum* shows an average mating rate of about 6-15 copulations per hour (Schärer et al. 2004a; Janicke and Schärer 2009b; Marie-Orleach et al. 2013) and the high mating rate may lead to suck behaviour because individuals prefer donating sperm more than receiving (Schärer et al. 2004a; Vizoso et al. 2010). The worms are kept in cultures in glass petri dishes filled with artificial sea water (ASW, 32‰) or nutrient-enriched artificial seawater (Guillard’s f/2 medium) (Guillard and Ryther 1962) and fed with diatoms (*Nitzschia curvilineata*). They are kept under standard conditions on a 14:10 light:dark cycle at 60% relative humidity and a constant temperature of 20°C. All the animals used in this experiment as knockdown/control donors and as recipients (see below) belonged to the highly inbred DV1 line (Janicke et al. 2013) that was previously used to identify and functionally characterize putative seminal fluid candidates (Weber et al. 2018; Weber et al., submitted; Patlar et al., submitted).

For the identification of transcripts that are genetically correlated with mating frequency, we analysed mating behavior recorded in an earlier experiment by Patlar et al. 2019 (submitted). Briefly, the worms used in this experiment originated from 12 highly inbred lines (hereafter genotypes) that belong to a larger set of inbred lines which was originally generated at the University of Innsbruck and is now maintained at the University of Basel (for details, see Vellnow et al. 2017). Inbred lines are called DV lines, namely DV1 (Janicke et al. 2013; Wasik et al. 2015), DV3, DV8, DV13, DV28, DV69 and DV71 (Sekii et al. 2013) and DV47, DV49, DV61, DV75 and DV84 (Vellnow et al. 2017).

To assign paternity to offspring of competing ejaculate donors (i.e., what would be competing males in separate-sexed animals), we used an outbred transgenic BAS1 line of *M. lignano* that expresses GFP ubiquitously (Marie-Orleach et al. 2016; Vellnow et al. 2018) as sperm competitors. Thereby the resulting offspring could be unambiguously assigned as being sired by either the DV1 (GFP^−^) or BAS1 (GFP^+^) worm (see also Janicke et al. (2013) and Marie-Orleach et al. (2014), which employed a GFP-expressing inbred line (HUB1) for the same purpose; the BAS1 culture was created by backcrossing the HUB1 onto an outbred background). Offspring production, mating frequency, and morphology were previously found not to differ between individuals from HUB1 (GFP^+^) and DV1 (GFP^−^) lines (Marie-Orleach et al. 2014).

### Selection of candidate transcripts

To identify candidate transcripts for subsequent RNAi, we re-used data on SFP transcript expression reported in Patlar et al. (2018) together with novel data on mating frequency collected but not reported as part of that earlier experiment (Patlar et al. 2018). Briefly, 12 inbred lines of *M. lignano* were raised in two different social group sizes, namely groups of two or eight individuals. Social group size is a good predictor of mating group size and thus sperm competition intensity in *M. lignano* (Schärer and Ladurner 2003; Schärer et al. 2004b; Janicke and Schärer 2009a). All genotype/group size combinations were raised under controlled condition for approximately 7 weeks until behavioral and SFP expression measurements were carried out (see details Patlar et al. 2018).

To examine the mating frequency, mating pairs where formed with one partner always coming from a single inbred line (DV1) to be used as the virgin, standardized recipient grown under strict isolated conditions, and the other partner, the donor, originated from one of the different genotype/group size combinations. In total, 18 independent pair replicates and 18 independent octet replicates of each genotype were coupled with standardized recipients to observe mating behaviours. To be able to distinguish worms in mating pairs under normal light during observations, the recipient worms were exposed to the food coloring dye Grand Bleu [E131, E151] (Les Artistes – Paris), diluted to a concentration of 0.25 mg/mL in ASW, for 24 h beforehand. Such a 24 h exposure enables us to easily distinguish colored from non-colored worms, and has previously been shown not to affect the mating frequency (Marie-Orleach et al. 2013). Observations were done by video recording, each donor worm and its colored recipient partner was transferred into two-dimensional, mating observation chambers described in detail elsewhere (Schärer et al. 2004a). Briefly, the recipient worm was transferred with a 1.7 μl drop of ASW to the center of a coated (Sigmacote®, Saint Louis, USA) microscope slide which has four HERMA photo stickers (Filderstadt, Germany) as spacers adhered to the sides (two per side). Then the donor worm was transferred with 1.7 μl ASW to the recipient, merging the two drops. Each observation chamber consisted of 16 such pairs, each made by 3.4 μl drops of ASW, on a single glass slide. Once all mating pairs had been placed on the slide, another coated microscope slide was placed upon the first, to form each drop into a shallow three-dimensional pool that the worms can swim inside (Fig. S1). To protect the drops from evaporation, the area was sealed with a thin line of Vaseline around the perimeter prior to adding the second slide. Immediately after the observation chamber was ready, it was placed under a camera for a video recording of 2 hours. Video recordings were obtained with a DFK 41AF02 camera (Imaging source GmbH, Bremen, Germany) connected to a computer running the software Debut Video Capture Professional, version 2.02. The videos were captured at one frame per second, with a frame rate of ten (a video of 2 hours was zipped into 12 mins), in .mov format with 1920×1080 HD resolution. Videos were analyzed using the Kinovea video player, version 0.8.15. For analysis of mating frequency, the number of copulations for each mating pair was counted throughout the full 2h period based on the videos recorded.

We then identified candidates in two steps. First, we used the first four principal components (PCs) describing variation in seminal fluid transcript expression from Patlar et al. (2018) to estimate which (if any) of these were genetically correlated with mating frequency, in both pairs and octets. Based on identifying one such PC, namely PC4, which was consistently negatively correlated with mating frequency at both social group sizes, we then investigated further those transcripts that were significantly loaded on PC4. We directly assessed how SFP expression level correlated with mating frequency, predicting that candidate mediators of a reduced mating frequency in partners should exhibit a negative genetic correlation between expression level and mating frequency. Two such candidates (Mlig-pro46 and Mlig-pro63, see Results) were then investigated further in an RNAi knockdown experiment.

Because the same two candidate transcripts, Mlig-pro46 and Mlig-pro63, were also positively loaded on an additional axis of seminal fluid variation, PC3, and this was positively correlated with suck propensity in the pairs environment (Patlar et al, submitted), we further investigated whether RNAi knockdown also impacted on partner suck behavior.

### Raising conditions for the RNAi knockdown experiment

6 to 8 days post-hatching, a batch of same-age hatchlings (to be used as donors and recipients) was collected and distributed in glass petri dishes filled with ASW and fed *ad libitum* with algae at a density of ~150 individuals per dish. Individuals were transferred once per week to new glass petri dishes filled with ASW and *ad libitum* algae until they underwent tail amputation (day 40 and 36 for the doors and recipients respectively – see below).

### RNA interference

RNAi was performed as previously described (Kuales et al. 2011), largely following the same procedures as in Weber et al. (submitted) to ensure comparability of results. Briefly, for both seminal fluid candidates, a double-stranded RNA (dsRNA) probe was generated by an *in vitro* transcription system using primer pairs with T7 and SP6 promoter regions (T7 and SP6 Ribomax™ large scale RNA kit, Promega) (Mlig-pro46: forward primer: CTGCACGGTTGTTACCTTCG, reverse primer: TCATCTTCATAATTGCGGTGAAAG; Mlig-pro63: forward primer: ACAACTGACAATGCGATTAGC, reverse primer: CTGCTCGTACACAACCATCG). In the control treatment we added just water instead of dsRNA; control individuals were otherwise treated identically to the knockdown individuals. Previous studies with dsRNA for firefly luciferase indicate receipt of dsRNA per se does not affect worms (Pfister et al. 2008; Sekii et al. 2009; Arbore et al. 2015; Lengerer et al. 2018; Weber et al. submitted), so here we followed several other *M. lignano* studies in using a no dsRNA probe as the negative control (Kuales et al. 2011; Lengerer et al. 2014; Ramm et al. 2019). The efficacy of RNAi knockdown was verified by performing whole mount in situ hybridization (ISH) in a preliminary test (Fig. S1).

Before beginning the RNAi (or control) treatment, donor animals were tail-amputated between the antrum and ovaries. This takes advantage of the regenerative capacity of *M. lignano* (Egger et al. 2006), allowing us to remove the antrum, with all potential previously received ejaculate in it, the seminal vesicle, with potential (own) stored sperm as well as the SFP-producing prostate gland cells. Through amputation and subsequent regeneration, we ensured that seminal fluid production was ‘reset’ prior to the RNAi/control treatment and the individuals have an equal amount and age of stored sperm and seminal fluid reserves and further that individuals also contained no received sperm or seminal fluid at the beginning of the mating trials (see below). After amputation, individuals were randomly allocated to one of four treatment groups: either the negative (no dsRNA) control or one of three RNAi treatments containing the relevant dsRNA solution (Mlig-pro46 only; Mlig-pro63 only; or Mlig-pro46 and Mlig-pro63 combined). They were maintained in these treatment groups throughout the regeneration process, kept individually in a well of a 60-well microtest plate (Greiner Bio-One™ 60-well HLA Terasaki Plates). Each worm was placed in 10μl dsRNA solution (~25 ng/μl dsRNA for the specific transcript (in combined treatment in total 50ng/μl dsRNA) in ASW-algae mix). Throughout the whole experiment, animals were fed *ad libitum* with algae and were maintained under standard culture conditions. On day 41, 43, 45, 47 and 49, 2μl of the relevant dsRNA solution was added to each RNAi well. On day 42, 44, 46, and 48, the worms were transferred to a new well containing 10μl of new dsRNA solution to ensure a constant exposure to dsRNA. The first mating trial was conducted on day 10 post-amputation, which is sufficient time to allow complete regeneration (Egger et al. 2006; Lengerer et al. 2018).

The GFP^+^ donor worms used in the experiment as sperm competitors to the experimental subjects were also tail-amputated on the same day as the knockdown/control worms at the age of 40 (+/− 1) days. Thereafter, they were also each kept individually in one well of a 60-well microtest plate in 10μl ASW with *ad libitum* algae. GFP^+^ worms were transferred to a new well containing 10μl ASW with *ad libitum* algae once on day 45.

The recipient worms used in the experiment were 50 (+/− 1) day old adult worms, and were also tail-amputated, 14 days prior the mating assay at age 36 (+/− 1) days. After amputation, they were also each kept individually in one well of a 60-well microtest plate in 10μl ASW with *ad libitum* algae. Recipient worms were transferred to a new well containing 10μl ASW with *ad libitum* algae once on day 43.

### Observation of mating and suck frequency of knockdown

For each RNAi/control treatment, we then conducted two separate assays with separate batches of donor, competitor and recipient worms, to measure (i) mating frequency (in both batches), (ii) suck propensity (in both batches), and (iii) either defensive (*P*_1_) or offensive (*P*_2_) sperm competitive ability (depending on the batch: in batch 1 we measured only *P*_1_ and in batch 2 only *P*_2_). Initially, worms in both assays (*P*_1_ & *P*_2_) were treated identically. The mating frequency and suck propensity were calculated by counting the number of copulations and the number of suck events per mating by the recipient during the 2.5 hours mating period which was filmed during the sperm competitive ability assays described in the next subsection. Video recording was performed using mating chambers using the same procedure as described above, with the only exception being that the mating period lasted for 2.5 hours here instead of the 2 hours used for the earlier trials identifying candidates.

### Sperm competitive ability assays (*P*_1_ & *P*_2_)

All mating trials were conducted on day 11 post-amputation. Recipients were kept after their mating trial in 60-well plates in ASW with *ad libitum* algae and were transferred to a new well every second day until day 11 (6 wells in total), where they remained until day 21 (after which no further offspring were detected). The resulting offspring were counted and categorized as either GFP^−^ (sired by first knockdown or control donor) or GFP^+^ (sired by the competitor donor) until day 21, based on expression of GFP assessed at age 7-10 days using a Nikon SMZ-18 stereomicroscope with a C-HGFI Intensilight fluorescence light source and GFP filter cube (Nikon GmbH, Düsseldorf, Germany).

To estimate defensive sperm competitive ability (*P*_1_), either knockdown or control worms were mated and filmed for 2.5 hours with a randomly selected recipient worm in the above-described mating observation chamber. After 2.5 hours the recipient worms were each put individually in a well of a 60-well microtest plate. 30 mins after separating the mating partners, a GFP^+^ sperm competitor worm was added to the well containing the already-mated recipient worm, and the pair was allowed to mate for a further 2.5 hours. After the 2.5 hours mating period, the recipient and the GFP^+^ sperm competitor were separated into an individual well as described above.

To estimate offensive sperm competitive ability (*P*_2_), the sperm competition assay was carried out exactly like the *P*_1_ assay, except that the GFP^+^ worm was paired with the recipient first and the knockdown/control worm second. Again, the mating period with the knockdown/control individual was conducted in the mating observation chamber and filmed.

Each treatment group started with 36 donor worms at the beginning of the RNAi treatment. With some replicates excluded due to death of the donor before the mating trial, the absence of resulting offspring or the observation of five or fewer successful matings, the final realized sample sizes for each treatment group ranged from 22-34 (see Table 1).

**Table 1.** Descriptive statistics and tests for treatment effects on mating rate, suck propensity and sperm competitive ability following RNAi knockdown of 2 seminal fluid transcripts (Mlig-pro46; Mlig-pro63; Mlig-pro46 & Mlig-pro63 combined).

| | $P_1$ | | | | | | $P_2$ | | | | | |
| --- | --- | --- | --- | --- | --- | --- | --- | --- | --- | --- | --- | --- |
| Treatment | n | Mean paternity share ($P_1$) | Std. Error | df | t-value | p-value | n | Mean paternity share ($P_2$) | Std. Error | df | t-value | p-value |
| control | 32 | 0.18 | 0.05 |  |  |  | 27 | 0.88 | 0.05 |  |  |  |
| Mlig-pro46 | 28 | 0.20 | 0.05 | 58 | 1.01 | 0.32 | 22 | 0.92 | 0.03 | 47 | 0.77 | 0.44 |
| Mlig-pro63 | 30 | 0.21 | 0.07 | 60 | 0.37 | 0.71 | 25 | 0.90 | 0.04 | 50 | 0.44 | 0.66 |
| combined | 30 | 0.23 | 0.05 | 60 | 0.62 | 0.54 | 26 | 0.86 | 0.05 | 51 | 0.37 | 0.71 |
|  | n | Mean number of matings | Std. Error | df | t-value | p-value | n | Mean number of matings | Std. Error | df | t-value | p-value |
| control | 33 | 20.06 | 1.74 |  |  |  | 30 | 15.27 | 1.13 |  |  |  |
| Mlig-pro46 | 31 | 21.13 | 1.43 | 62 | 0.48 | 0.63 | 23 | 16.74 | 1.35 | 51 | 0.85 | 0.40 |
| Mlig-pro63 | 34 | 22.03 | 1.44 | 65 | 0.87 | 0.39 | 28 | 17.11 | 1.34 | 56 | 1.02 | 0.31 |
| combined | 32 | 23.06 | 1.41 | 63 | 1.37 | 0.18 | 28 | 16.29 | 1.40 | 56 | 0.57 | 0.57 |
|  | n | Mean suck propensity | Std. Error | df | t-value | p-value | n | Mean suck propensity | Std. Error | df | t-value | p-value |
| control | 33 | 0.54 | 0.02 |  |  |  | 29 | 0.47 | 0.03 |  |  |  |
| Mlig-pro46 | 31 | 0.51 | 0.02 | 62 | -1.135 | 0.26 | 24 | 0.44 | 0.04 | 51 | -0.768 | 0.45 |
| Mlig-pro63 | 34 | 0.44 | 0.03 | 65 | -3.032 | 0.0035 | 26 | 0.41 | 0.03 | 53 | -1.543 | 0.1290 |
| combined | 32 | 0.47 | 0.03 | 63 | -1.821 | 0.07 | 27 | 0.41 | 0.03 | 54 | -1.758 | 0.08 |

### Statistical analysis

The effect of genotype on mating frequency was assessed by performing two-way ANOVA with interaction after testing for homogeneity of variance for mating group size by performing Levene tests. Genetic correlation estimates were based on the Pearson correlation coefficients calculated between mean trait values of genotypes. We calculated the corresponding genetic correlations as r_G_ = Cov(x1, x2) / [Var(x1) × Var(x2)], and tested for statistical significance by comparing the z-scores to two-tailed significance levels derived from a standard normal distribution.

For analysis of the *P*_1_ and *P*_2_ assays, the paternity share of knockdown and control individuals (GFP^−^) were compared against the GFP^+^ competitor using a generalized linear model with a quasibinomial distribution and a logit link function (Engqvist 2013). For analysis of the mating frequency, we compared the number of copulations of knockdown and control mating pairs, using a separate linear model for each of the three treatment-control comparisons. For analysis of suck propensity, the frequency of sucking events (i.e. the proportion of matings followed by a suck) of the mating partners of knockdown and control individuals were compared using a generalized linear model with a binomial distribution and a logit link function. Analyses were conducted using the lme4 package (Bates et al. 2015) in R (R version 3.1.3 2015).

## Results

### Genetic correlations between SFP expression, mating frequency and suck propensity

We estimated genetic correlations between mating frequency and the overall seminal fluid investment (PC1) and relative composition (PC2-4) axes reported in Patlar et al. (2018) at two different group sizes (pairs and octets). We found that mating frequency was not genetically correlated with overall seminal fluid investment, but there was a highly significant negative genetic correlation between mating frequency and PC4 in both pairs and octets (Fig. 1a).

**Fig. 1.**
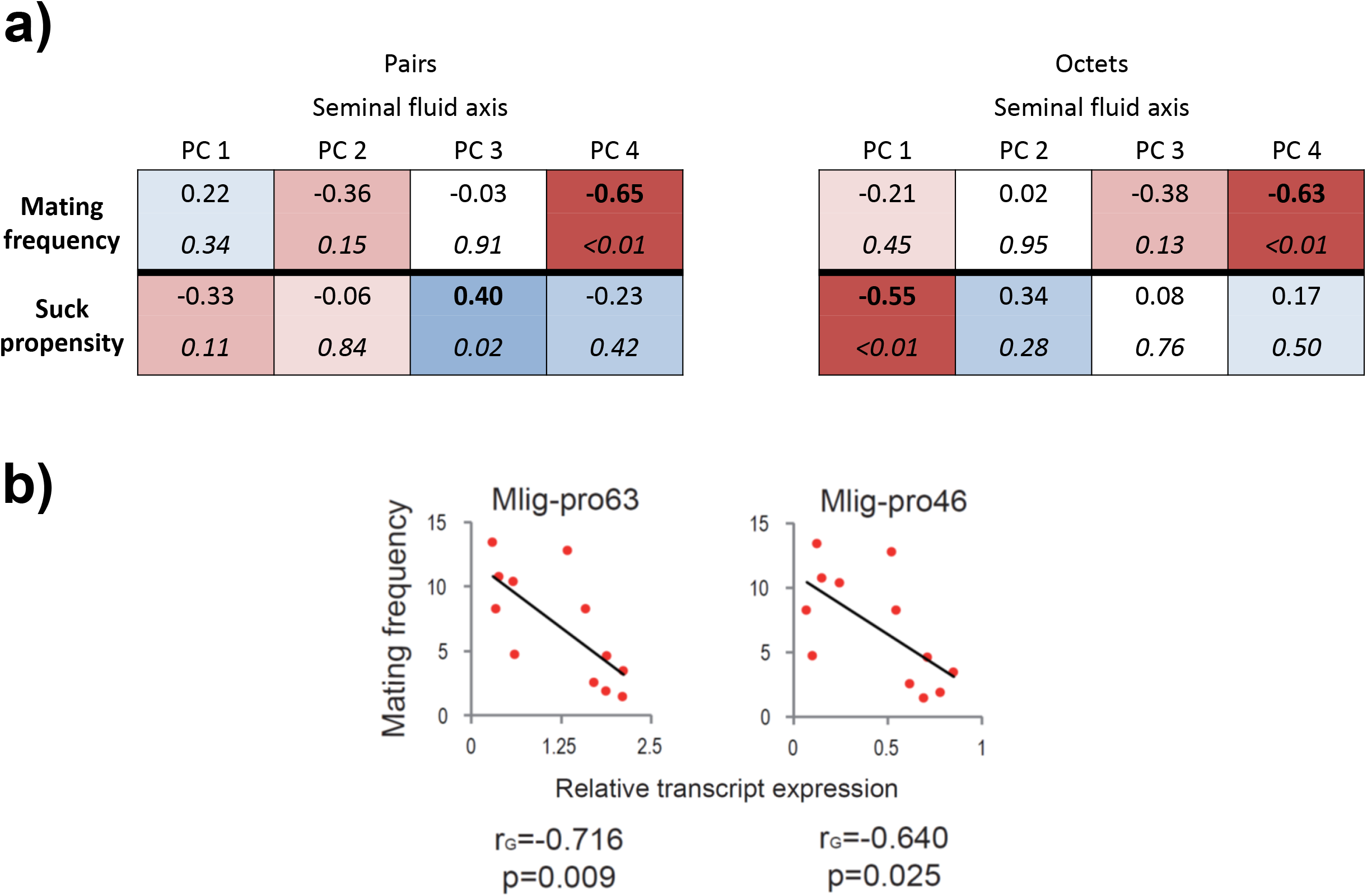
Genetic correlations (r_G_) between seminal fluid of the donor and mating rate. The transcripts are labelled according to their Mlig-pro[number] identifier assigned in Weber et al. (2018). a) Genetic correlation coefficients and P-values (italic) for seminal fluid axes (principal components) and mating rate/suck propensity in pairs and octets. b) The relationship between the average mating rate and relative transcript expression with genetic correlation coefficients and respective p-values in octets.

Having established that mating frequency is highly negatively correlated with PC4, we therefore next estimated the genetic correlations between mating frequency and the five seminal fluid transcripts which were significantly loaded on PC4 (Patlar et al. 2018). Among these, we found three transcripts which exhibit a significant negative correlation between mating frequency and SFP transcript expression: Mlig-pro63, Mlig-pro46 and Mlig-pro37 (Fig. 1b; here we illustrated these correlations only for octets, which is anyway the more relevant environment for sperm competition, because both group sizes show a very similar pattern). In fact, according to their highly similar sequence with overlapping regions, and the fact that when blasted against the *M. lignano* genome assembly ML2 (Wasik et al. 2015) they align to the same regions within the same protein coding gene in the genome, Mlig-pro37 and Mlig-pro46 appear to belong to the same gene. For that reason, for our RNAi screen we selected just the transcript Mlig-pro46, which was already investigated in the previous screen (Weber et al., submitted), plus the independent candidate Mlig-pro63.

Notably, these same candidates were also significantly loaded on PC3, as reported before in Patlar et al. 2018, which was found as to be positively correlated with suck behavior (Fig. 1a), at least in the pair group size (Patlar et al submitted).

### RNAi knockdown effects on mating frequency

When we compared the mating frequency of mating pairs including an SFP knockdown donor to those including a control donor, none of the knockdowns impacted strongly on mating frequency, as measured by the total number of copulation events in the 2.5 hour mating period (all p ≥ 0.1, Fig. 2; for full statistical details for each knockdown, see Table 1).

**Fig. 2.**
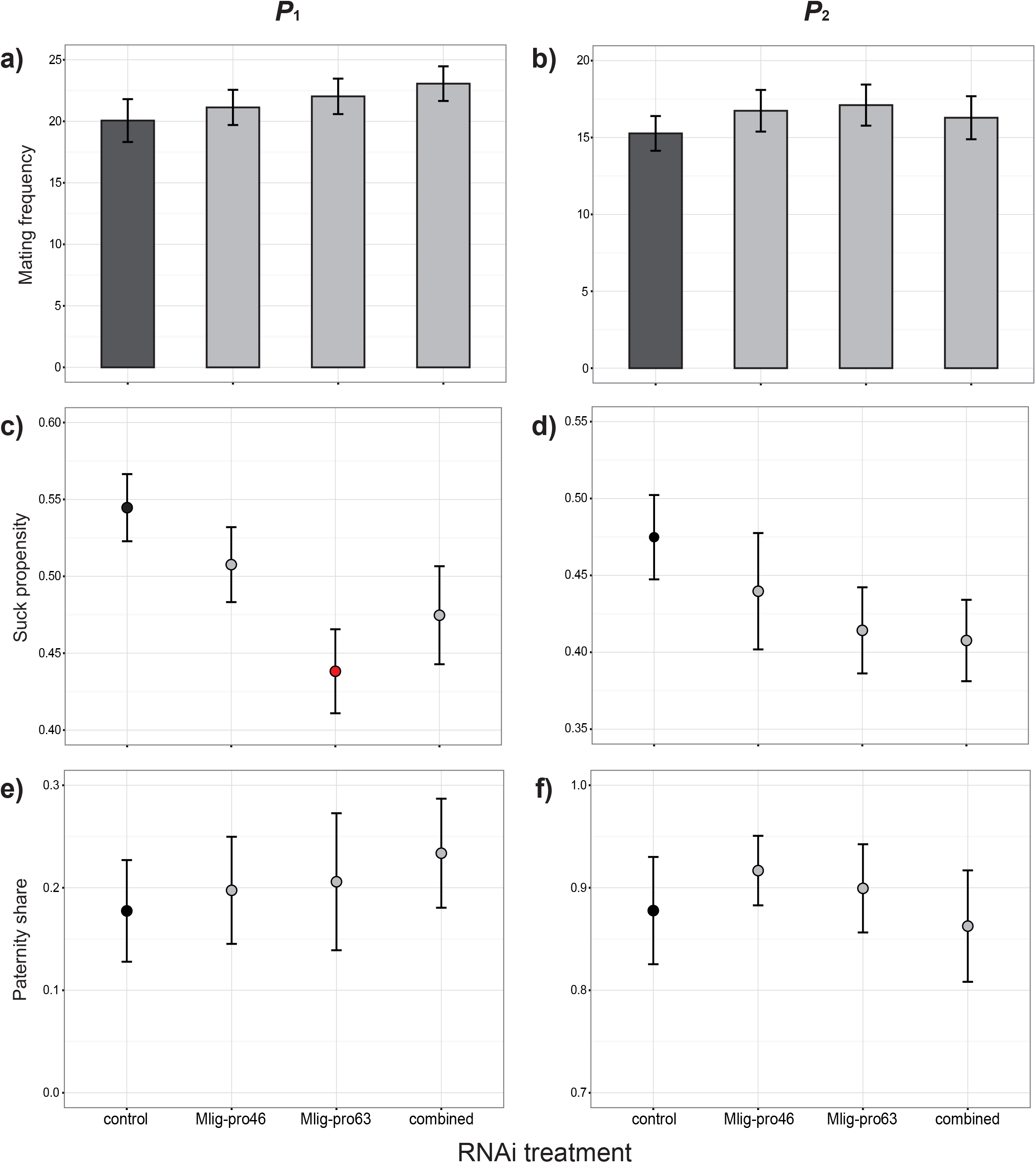
The effect of RNAi knockdown of 2 different seminal fluid transcripts (Mlig-pro46; Mlig-pro63; Mlig-pro46 & Mlig-pro63 combined) The transcripts are labelled according to their Mlig-pro[number] identifier assigned in Weber et al. (2018). a) Mean mating rate ±SE of knockdown versus control mating pairs when the RNAi worm mated first. b) Mean mating rate ±SE of knockdown versus control mating pairs when the RNAi worm mated second. c) Mean suck propensity ±SE of the ejaculate receiving individual of knockdown versus control mating pairs when the RNAi worm mated first. d) Mean suck propensity ±SE of the ejaculate receiving individual of knockdown versus control mating pairs when the RNAi worm mated second. e) Mean paternity share (*P*_1_) ±SE of knockdown versus control individuals mated with a partners when the RNAi worm mated first. f) Mean paternity share (*P*_2_) ±SE of knockdown versus control individuals mated with a partners when the RNAi worm mated second.

### RNAi knockdown effects on suck propensity

When we compared the mean suck propensity of mating pairs including a SFP knockdown donor to those including a control donor, one of the individual knockdowns exhibited a strongly reduced suck propensity (Mlig-pro63, t = −3.032, p = 0.0035; Fig. 2; Table 1), as measured by the sucking events per mating in the 2.5 hour mating period, when they mated as the first partner with a recipient worm who subsequently mated with an outbred sperm competitor. None of the other knockdowns had a significant effect on suck propensity, though in both the *P*_1_ and *P*_2_ assays the combined RNAi showed a non-significant trend in the same direction for suck propensity.

### RNAi knockdown effects on *P*_1_ and *P*_2_

When we compared recipient worms mated to SFP knockdown donors to those mated to the control donors, none of the knockdowns impacted strongly on defensive sperm competitive ability (*P*_1_) or offensive sperm competitive ability (*P*_2_), as measured by the paternity share between knockdown/control individuals and the competitor (all p ≥ 0.3, Fig. 2; Table 1).

## Discussion

By selecting two putative seminal fluid transcripts with prostate-limited expression (Weber et al. 2018) that exhibit genetic correlations with both mating frequency and partner suck propensity (Patlar et al. 2018; Patlar submitted), and subjecting these to RNAi knockdown (individually and combined) followed by behavioral and competitive paternity assays, we aimed to test whether these proteins directly influence partner behavior. In fact, we did not detect any significant seminal fluid-mediated effect on mating frequency. On the other hand, we did find that one of the seminal fluid transcripts, Mlig-pro63, appears to impact positively on the frequency of the post-copulatory suck behavior often exhibited by ejaculate recipients in *M. lignano*. Finally, we also did not detect any difference in paternity share between knockdown and control individuals, neither when the knockdown individuals were the first mating partners (*P*_1_) or when the knockdown individuals were the second mating partners (*P*_2_) in controlled sperm competition assays. In the following, we discuss each of these three main results in turn.

Firstly, one reason why RNAi knockdown of Mlig-pro46 and/or Mlig-pro63 had no detectable impact on mating frequency could be that the SFPs tested in our assay have a long-term effect, and not a more or less immediate one as we tested for here, on subsequent behaviors like mating frequency. In our experimental design, we could only detect possible effects between the first copulation and the end of the 2.5 hours recording time. While we thus checked for immediate effects on re-mating, there are for example SFPs in *Drosophila* known to act only over the longer term. In a screen of 25 *D. melanogaster* SFP knockdowns, none appeared to modulate the receptivity of the mated female at 24 h post mating (Ram and Wolfner 2007b), with an equally low receptivity to re-mating regardless of whether females mated to control or knockdown males. But three of these SFP knockdowns showed a significant long-term effect on female receptivity, with females mated to males from these three knockdown treatments being significantly more receptive to re-mating at 4 d post-mating than were mates of control males. Nevertheless, the way that we identified candidates in *M. lignano* involved testing for correlations between transcript expression and mating frequency over a similarly short time-scale, so the lack of effect in our assay is still surprising. Perhaps another explanation could be that while Patlar et al. (Patlar et al. 2018; Patlar unpublished) used 12 different inbred lines, differing in their genotypes, we used in our experiment only one genotype. The 12 inbred lines could differ in other, correlated ways which in combination with Mlig-pro46 and/or Mlig-pro63 affect mating frequency, but are not incorporated in our RNAi experimental design.

Our results indicate that Mlig-pro63 appears to play a role in promoting the suck behavior of the mating partner. That there is a decrease of the suck propensity with the loss of this protein from the ejaculate is at first glance also a surprising result, at least from a donor perspective. Indeed, Patlar et al. (submitted) have recently identified two SFPs in *M. lignano* that have the opposite effect, apparently manipulating partner behavior to reduce the frequency of sucking and therefore presumably gaining greater control over the fate of the transferred ejaculate. Consistent with this, previous work had indicated that individuals mated to virgin partners (which presumably transfer bigger ejaculates) exhibit a lower frequency of the suck behavior (Marie-Orleach et al. 2013).

By contrast, the effect we detected – of a decrease in suck propensity upon the RNAi-induced loss of one ejaculate component – suggests that other SFPs might act as cues used by the ejaculate recipient to trigger suck behavior. This would seem to imply that they have evolved and are still included in the ejaculate for some other, as yet unknown, reason that still provides a net benefit to the ejaculate donor, since – assuming the suck behavior benefits recipients at the expense of donors – their inclusion would otherwise seem to be a maladaptive strategy.

Also with respect to suck propensity, we note that the fact we observed an effect of Mlig-pro63 knockdown in the *P*_1_ but not the *P*_2_ assay suggests that Mlig-pro63 presumably transferred first by the sperm competitor individual in the *P*_2_ assay was still affecting partner behavior at the time of pairing with the Mlig-pro63 knockdown worm. This highlights one difficulty with performing such double mating assays, in that SFP-mediated effects presumably intended to influence the utilization of own sperm can, under certain study designs that may not well reflect the situation in nature, actually influence that of rivals, and vice versa (see e.g. Mueller et al. 2008; Nguyen and Moehring 2018; see also Hodgson and Hosken 2006).

The absence of detectable impacts of SFP knockdown on sperm competitiveness could also be a direct result of the fact the time window we investigated does not well reflect the action of these proteins. Also, because in our experiment the competitor was introduced either very soon following the knockdown/control individuals (*P*_1_ assay), or else there was no competitor following the knockdown/control individuals (*P*_2_ assay), there was perhaps no opportunity to influence sperm competition outcomes via (eventually) reduced receptivity. The surprising result that we did not, contrary to the previous RNAi screen (Weber et al., submitted), find an effect of the Mlig-pro46 knock-down on paternity share, certainly suggests we should remain cautious about interpreting the role of Mlig-pro46 in sperm competition until we have gained a greater understanding of its mechanism of action. However, the different outcomes of the two studies could perhaps be due to the slightly different experimental designs regarding mating duration and length of the break between the two competitors. While in the previous experiment there was one hour between removing the first donor and adding the second, as well as a longer mating period of 3 instead of 2.5h, in this experiment this gap between pairings was just half an hour. The shorter mating period presumably reduced the opportunity for cumulative effects on sperm competitiveness, and the shorter gap is likely to have exacerbated any carry-over effects alluded to above, meaning effects of SFPs presumably intended for own ejaculates actually also impact on rival ejaculates, or vice versa, tending to equalize paternity success between competitors.

In conclusion, we found evidence for seminal fluid-mediated effects on suck propensity in the simultaneously hermaphrodite *M. lignano*, but no indication that the two candidate proteins Mlig-pro46 and Mlig-pro63 affect mating frequency. Further research will be needed to investigate a potential impact of seminal fluid on the long-term receptivity of mating partners in *M. lignano*. Overall, by using a combination of quantitative genetic and behavioral data to first identify seminal fluid components with potential effects on post-mating behavior and subsequent RNAi knockdown assays, we have gained some novel insights about seminal fluid action in the flatworm *M. lignano*, and a greater appreciation of the central importance of the suck behavior to its reproductive interactions. The combination of quantitative genetics and behavioral data together with subsequent RNAi could be a promising approach for future investigation of SFP function in this and other non-model organisms.

**Fig. S.1 – Representative whole-mount in situ hybridization expression patterns for knockdown of the 2 different seminal fluid transcripts (Mlig-pro46; Mlig-pro63; Mlig-pro46 & Mlig-pro63 combined)**. ISH patterns for control individuals (green) compared to knockdown individuals (blue) for the respective transcript.

## References

Abraham, S., L. A. Lara-Pérez, C. Rodríguez, Y. Contreras-Navarro, N. Nuñez-Beverido, S. Ovruski, and D. Pérez-Staples. 2016. The male ejaculate as inhibitor of female remating in two tephritid flies. J. Insect Physiol. 88:40–47.

Arbore, R., K. Sekii, C. Beisel, P. Ladurner, E. Berezikov, and L. Schärer. 2015. Positional RNA-Seq identifies candidate genes for phenotypic engineering of sexual traits. Front. Zool. 12:14.

Arnqvist, G., and L. Rowe. 2005. Sexual Conflict. Princeton University Press; Princeton.

Avila, F. W., L. K. Sirot, B. A. Laflamme, C. D. Rubinstein, and M. F. Wolfner. 2011. Insect Seminal Fluid Proteins : Identification and Function. Annu. Rev. Entomol. 56:21–40.

Bates, D., M. Maechler, B. Bolker, and S. Walker. 2015. Fitting linear mixed-effects models using lme4. J. Stat. Softw. 67:1–48.

Birkhead, T. R. 1995. Sperm competition: Evolutionary causes and consequences. Reprod. Fertil. Dev. 7:755–775.

Birkhead, T. R., and T. Pizzari. 2002. Postcopulatory sexual selection. Nat. Rev. Genet. 3:262–273.

Cameron, E., T. Day, and L. Rowe. 2007. Sperm Competition and the Evolution of Ejaculate Composition. Am. Nat. 169:E158–E172.

Chapman, T. 2001. Seminal fluid-mediated fitness traits in Drosophila. Heredity 87:511–521.

Chapman, T. 2018. Sexual conflict: mechanisms and emerging themes in resistance biology. Am. Nat. 192:217–229.

Chapman, T., J. Bangham, G. Vinti, B. Seifried, O. Lung, M. F. Wolfner, H. K. Smith, and L. Partridge. 2003. The sex peptide of *Drosophila melanogaster*: Female post-mating responses analyzed by using RNA interference. Proc. Natl. Acad. Sci. USA 100:9923–9928.

Chapman, T., L. F. Liddle, J. M. Kalb, M. F. Wolfner, and L. Partridge. 1995a. Cost of mating in *Drosophila melanogaster* females is mediated by male accessory gland products. Nature, doi: 10.1038/373241a0.

Chapman, T., L. F. Liddle, J. M. Kalb, M. F. Wolfner, L. Partridge, T. Chapman, L. Partridge, J. M. Kalb, and M. F. Wolfner. 1995b. Cost of mating in *Drosophila melanogaster* females is mediated by male accessory gland products. Nature 373:241–244.

Chapman, T., D. M. Neubaum, M. F. Wolfner, and L. Partridge. 2000. The role of male accessory gland protein Acp36DE in sperm competition in *Drosophila melanogaster*. Proc. R. Soc. B-Biological Sci. 267:1097–1105.

Charnov, E. L. 1979. Simultaneous hermaphroditism and sexual selection. Proc. Natl. Acad. Sci. USA 76:2480–2484.

Eberhard, W. G. 1996. Female Control: Sexual Selection by Cryptic Female Choice. Princeton University Press.

Egger, B., P. Ladurner, K. Nimeth, R. Gschwentner, and R. Rieger. 2006. The regeneration capacity of the flatworm *Macrostomum lignano* - On repeated regeneration, rejuvenation, and the minimal size needed for regeneration. Dev. Genes Evol. 216:565–577.

Engqvist, L. 2013. A general description of additive and nonadditive elements of sperm competitiveness and their relation to male fertilization success. Evolution 67:1396–1405.

Forsberg, G., I. Bednar, P. Eneroth, and P. Södersten. 1990. β-Endorphin acts on the reproductive tract of female rats to suppress sexual receptivity. Neurosci. Lett. 115:92–96.

Gillott, C. 2003. Male accessory gland secretions: modulators of female reproductive physiology and behavior. Annu. Rev. Entomol. 48:163–184.

Guillard, R. R. L., and J. H. Ryther. 1962. Studies of marine planktonic diatoms. I. *Cyclotella nana* Hustedt, and *Detonula confervacea* (cleve) Gran. Can. J. Microbiol. 8:229–239.

Harmer, A. M. T., P. Radhakrishnan, and P. W. Taylor. 2006. Remating inhibition in female Queensland fruit flies: Effects and correlates of sperm storage. J. Insect Physiol. 52:179–186.

Harshman, L. G., and T. Prout. 1994. Sperm displacement without sperm transfer in *Drosophila melanogaster*. Evolution 48:758–766.

Häsemeyer, M., N. Yapici, U. Heberlein, and B. J. Dickson. 2009. Sensory neurons in the *Drosophila* genital tract regulate female reproductive behavior. Neuron 61:511–518.

Hodgson, D. J., and D. J. Hosken. 2006. Sperm competition promotes the exploitation of rival ejaculates. J. Theor. Biol. 243:230–234.

Hollis, B., M. Koppik, K. U. Wensing, H. Ruhmann, E. Genzoni, B. Erkosar, T. J. Kawecki, C. Fricke, and L. Keller. 2019. Sexual conflict drives male manipulation of female postmating responses in Drosophila melanogaster. Proc. Natl. Acad. Sci., doi: 10.1073/pnas.1821386116.

Hopkins, B. R., I. Sepil, and S. Wigby. 2017. Seminal fluid. Curr. Biol. 27:R404–R405.

Hyman, L. H. 1951. The Invertebrates: Platyhelminthes and Rhynchocoela. McGraw-Hill New York.

Janicke, T., L. Marie-Orleach, K. De Mulder, E. Berezikov, P. Ladurner, D. B. Vizoso, and L. Schärer. 2013. Sex allocation adjustment to mating group size in a simultaneous hermaphrodite. Evolution 67:3233–3242.

Janicke, T., and L. Schärer. 2009a. Determinants of mating and sperm-transfer success in a simultaneous hermaphrodite. J. Evol. Biol. 22:405–415.

Janicke, T., and L. Schärer. 2009b. Sex allocation predicts mating rate in a simultaneous hermaphrodite. Proc. R. Soc. B Biol. Sci., doi: 10.1098/rspb.2009.1336.

Kimura, K., K. Shibuya, and S. Chiba. 2013. The mucus of a land snail love-dart suppresses subsequent matings in darted individuals. Anim. Behav. 85:631–635.

Koene, J. M. 2006. Tales of two snails: Sexual selection and sexual conflict in *Lymnaea stagnalis* and *Helix aspersa*. Integr. Comp. Biol. 46:419–429.

Koene, J. M., W. Sloot, K. Montagne-Wajer, S. F. Cummins, B. M. Degnan, J. S. Smith, G. T. Nagle, and A. ter Maat. 2010. Male accessory gland protein reduces egg laying in a simultaneous hermaphrodite. PLoS One 5:e10117.

Kuales, G., K. De Mulder, J. Glashauser, W. Salvenmoser, S. Takashima, V. Hartenstein, E. Berezikov, W. Salzburger, and P. Ladurner. 2011. Boule-like genes regulate male and female gametogenesis in the flatworm *Macrostomum lignano*. Dev. Biol. 357:117–132.

Ladurner, P., L. Schärer, W. Salvenmoser, and R. M. Rieger. 2005. A new model organism among the lower Bilateria and the use of digital microscopy in taxonomy of meiobenthic Platyhelminthes: *Macrostomum lignano*, n. sp. (Rhabditophora, Macrostomorpha). J. Zool. Syst. Evol. Res. 43:114–126.

Lengerer, B., R. Pjeta, J. Wunderer, M. Rodrigues, R. Arbore, L. Schärer, E. Berezikov, M. W. Hess, K. Pfaller, B. Egger, S. Obwegeser, W. Salvenmoser, and P. Ladurner. 2014. Biological adhesion of the flatworm *Macrostomum lignano* relies on a duo-gland system and is mediated by a cell type-specific intermediate filament protein. Front. Zool. 11:12.

Lengerer, B., J. Wunderer, R. Pjeta, G. Carta, D. Kao, A. Aboobaker, C. Beisel, E. Berezikov, W. Salvenmoser, and P. Ladurner. 2018. Organ specific gene expression in the regenerating tail of Macrostomum lignano. Dev. Biol. 433:448–460.

Liu, H., and E. Kubli. 2003. Sex-peptide is the molecular basis of the sperm effect in Drosophila melanogaster. Proc. Natl. Acad. Sci. USA 100:9929–9933.

Marie-Orleach, L., T. Janicke, and L. Schärer. 2013. Effects of mating status on copulatory and postcopulatory behaviour in a simultaneous hermaphrodite. Anim. Behav. 85:453–461.

Marie-Orleach, L., T. Janicke, D. B. Vizoso, P. David, and L. Schärer. 2016. Quantifying episodes of sexual selection: Insights from a transparent worm with fluorescent sperm. Evolution 70:314–328.

Marie-Orleach, L., T. Janicke, D. B. Vizoso, M. Eichmann, and L. Schärer. 2014. Fluorescent sperm in a transparent worm: validation of a GFP marker to study sexual selection. BMC Evol. Biol. 14:148.

Michiels, N. K. 1998. Mating conflicts and sperm competition in simultaneous hermaphrodites. Pp. 219–254 *in* Sperm competition and sexual selection (eds Birkhead T., Møller A. P.). London, UK: Academic Press.

Moya-Laraño, J., and C. W. Fox. 2006. Ejaculate size, second male size, and moderate polyandry increase female fecundity in a seed beetle. Behav. Ecol. 17:940–946.

Mueller, J. L., J. R. Linklater, K. R. Ram, T. Chapman, and M. F. Wolfner. 2008. Targeted gene deletion and phenotypic analysis of the *Drosophila melanogaster* seminal fluid protease inhibitor Acp62F. Genetics 178:1605–1614.

Nakadera, Y., E. M. Swart, J. N. A. Hoffer, O. Den Boon, J. Ellers, and J. M. Koene. 2014. Receipt of seminal fluid proteins causes reduction of male investment in a simultaneous hermaphrodite. Curr. Biol. 24:859–862.

Neubaum, D. M., and M. F. Wolfner. 1999. Mated *Drosophila melanogaster* females require a seminal fluid protein, Acp36DE, to store sperm efficiently. Genetics 153:845–857.

Nguyen, T. T. X., and A. J. Moehring. 2018. A male’s seminal fluid increases later competitors’ productivity. J. Evol. Biol. 31:1572–1581.

Parker, G. A. 1979. Sexual selection and sexual conflict. Sex. Sel. Reprod. Compet. insects 123:166.

Parker, G. A. 1970. Sperm competition and its evolutionary consequences in the insects. Biol. Rev. 45:525–567.

Patlar, B., M. Weber, and S. A. Ramm. 2018. Genetic and environmental variation in transcriptional expression of seminal fluid proteins. Heredity 122:595–611.

Patlar, B., M. Weber, T. Temizyürek and S.A. Ramm n.d. Seminal fluid-mediated manipulation of post-mating behaviour in a simultaneous hermaphrodite. submitted.

Perry, J. C., and L. Rowe. 2008a. Ingested spermatophores accelerate reproduction and increase mating resistance but are not a source of sexual conflict. Anim. Behav. 76:993–1000.

Perry, J. C., and L. Rowe. 2008b. Neither mating rate nor spermatophore feeding influences longevity in a ladybird beetle. Ethology 114:504–511.

Pfister, D., K. De Mulder, V. Hartenstein, G. Kuales, G. Borgonie, F. Marx, J. Morris, and P. Ladurner. 2008. Flatworm stem cells and the germ line: Developmental and evolutionary implications of macvasa expression in *Macrostomum lignano*. Dev. Biol. 319:146–159.

Poiani, A. 2006. Complexity of seminal fluid: A review. Behav. Ecol. Sociobiol. 60:289–310.

Qazi, M. C. B. 2003. An early role for the *Drosophila melanogaster* male seminal protein Acp36DE in female sperm storage. J. Exp. Biol. 206:3521–3528.

R version 3.1.3. 2015. (2015-03-09) -- “Smooth Sidewalk” Copyright (C) 2015 The R Foundation for Statistical Computing Platform: x86_64-w64-mingw32/x64 (64-bit).

Radhakrishnan, P., and P. W. Taylor. 2008. Ability of male Queensland fruit flies to inhibit receptivity in multiple mates, and the associated recovery of accessory glands. J. Insect Physiol. 54:421–428.

Radhakrishnan, P., and P. W. Taylor. 2007. Seminal fluids mediate sexual inhibition and short copula duration in mated female Queensland fruit flies. J. Insect Physiol. 53:741–745.

Ram, K. R., and M. F. Wolfner. 2009. A network of interactions among seminal proteins underlies the long-term postmating response in *Drosophila*. Proc. Natl. Acad. Sci. USA 106:15384–15389.

Ram, K. R., and M. F. Wolfner. 2007a. Seminal influences: *Drosophila* Acps and the molecular interplay between males and females during reproduction. Integr. Comp. Biol. 47:427–445.

Ram, K. R., and M. F. Wolfner. 2007b. Sustained post-mating response in *Drosophila melanogaster* requires multiple seminal fluid proteins. Plos Genet. 3:2428–2438.

Ramm, S. A., B. Lengerer, R. Arbore, R. Pjeta, J. Wunderer, A. Giannakara, E. Berezikov, P. Ladurner, and L. Schärer. 2019. Sex allocation plasticity on a transcriptome scale: Socially sensitive gene expression in a simultaneous hermaphrodite. Mol. Ecol. 28:2321–2341.

Rönn, J., M. Katvala, and G. Arnqvist. 2006. The costs of mating and egg production in *Callosobruchus* seed beetles. Anim. Behav. 72:335–342.

Schärer, L. 2014. Evolution: Don’t be so butch, dear! Curr. Biol. 24:R311–R313.

Schärer, L., T. Janicke, and S. A. Ramm. 2015. Sexual conflict in hermaphrodites. Cold Spring Harb. Perspect. Biol. 7:a017673.

Schärer, L., G. Joss, and P. Sandner. 2004a. Mating behaviour of the marine turbellarian *Macrostomum* sp.: these worms suck. Mar. Biol. 145:373–380.

Schärer, L., and P. Ladurner. 2003. Phenotypically plastic adjustment of sex allocation in a simultaneous hermaphrodite. Proc. R. Soc. B Biol. Sci. 270:935–941.

Schärer, L., P. Ladurner, and R. M. Rieger. 2004b. Bigger testes do work more: Experimental evidence that testis size reflects testicular cell proliferation activity in the marine invertebrate, the free-living flatworm *Macrostomum sp*. Behav. Ecol. Sociobiol. 56:420–425.

Scharer, L., D. T. J. Littlewood, A. Waeschenbach, W. Yoshida, and D. B. Vizoso. 2011. Mating behavior and the evolution of sperm design. Proc. Natl. Acad. Sci. USA 108:1490–1495.

Schärer, L., and S. A. Ramm. 2016. Hermaphrodites. Pp. 212–224 *in* Encyclopedia of evolutionary biology (eds Kliman R). Oxford: Elsevier.

Sekii, K., W. Salvenmoser, K. De Mulder, L. Scharer, and P. Ladurner. 2009. Melav2, an elav-like gene, is essential for spermatid differentiation in the flatworm *Macrostomum lignano*. BMC Dev. Biol. 9:62.

Sekii, K., D. B. Vizoso, G. Kuales, K. De Mulder, P. Ladurner, and L. Schärer. 2013. Phenotypic engineering of sperm-production rate confirms evolutionary predictions of sperm competition theory. Proc. R. Soc. B Biol. Sci. 280:20122711.

Shutt, B., L. Stables, F. Aboagye-Antwi, J. Moran, and F. Tripet. 2010. Male accessory gland proteins induce female monogamy in anopheline mosquitoes. Med. Vet. Entomol. 24:91–94.

Simmons, L. 2001. Sperm Competition And Its Evolutionary Consequences In The Insects. Princeton University Press; Princeton.

Sirot, L. K., A. Wong, T. Chapman, and M. F. Wolfner. 2015. Sexual conflict and seminal fluid proteins: A dynamic landscape of sexual interactions. Cold Spring Harb. Perspect. Biol. 7:a017533.

Takami, Y., M. Sasabe, N. Nagata, and T. Sota. 2008. Dual function of seminal substances for mate guarding in a ground beetle. Behav. Ecol. 19:1173–1178.

Thornhill, R. 1983. Cryptic Female Choice and Its Implications in the Scorpionfly *Harpobittacus nigriceps*. Am. Nat. 122:765–788.

Uhl, G., S. H. Nessler, and J. M. Schneider. 2009. Securing paternity in spiders? A review on occurrence and effects of mating plugs and male genital mutilation. Genetica 138:75–104.

Vellnow, N., L. Marie-Orleach, K. S. Zadesenets, and L. Schärer. 2018. Bigger testes increase paternity in a simultaneous hermaphrodite, independently of the sperm competition level. J. Evol. Biol. 31:180–196.

Vellnow, N., D. B. Vizoso, G. Viktorin, and L. Schärer. 2017. No evidence for strong cytonuclear conflict over sex allocation in a simultaneously hermaphroditic flatworm. BMC Evol. Biol., doi: 10.1186/s12862-017-0952-9.

Vizoso, D. B., G. Rieger, and L. Schärer. 2010. Goings-on inside a worm: Functional hypotheses derived from sexual conflict thinking. Biol. J. Linn. Soc. 99:370–383.

Waage, J. K. 1986. Evidence for widespread sperm displacement ability among *Zygoptera* (Odonata) and the means for predicting its presence. Biol. J. Linn. Soc. 28:285–300.

Wasik, K., J. Gurtowski, X. Zhou, O. M. Ramos, M. J. Delás, G. Battistoni, O. El Demerdash, I. Falciatori, D. B. Vizoso, A. D. Smith, P. Ladurner, L. Schärer, W. R. McCombie, G. J. Hannon, and M. Schatz. 2015. Genome and transcriptome of the regeneration-competent flatworm, Macrostomum lignano. Proc. Natl. Acad. Sci. USA 112:12462–12467.

Weber, M., J. Wunderer, B. Lengerer, R. Pjeta, M. Rodrigues, L. Schärer, P. Ladurner, and S. A. Ramm. 2018. A targeted in situ hybridization screen identifies putative seminal fluid proteins in a simultaneously hermaphroditic flatworm. BMC Evol. Biol. 18:81.

Weber, M., A. Giannakara and S.A. Ramm n.d. Seminal fluid-mediated fitness effects in the simultaneously hermaphroditic flatworm *Macrostomum lignano*. submitted.

Wigby, S., and T. Chapman. 2005. Sex peptide causes mating costs in female *Drosophila melanogaster*. Curr. Biol. 15:316–321.

Yamane, T., T. Miyatake, and Y. Kimura. 2008. Female mating receptivity after injection of male-derived extracts in *Callosobruchus maculatus*. J. Insect Physiol. 54:1522–1527.

Yang, C. hui, S. Rumpf, Y. Xiang, M. D. Gordon, W. Song, L. Y. Jan, and Y. N. Jan. 2009. Control of the postmating behavioral switch in *Drosophila* females by internal sensory neurons. Neuron 61:519–526.

Yapici, N., Y. J. Kim, C. Ribeiro, and B. J. Dickson. 2008. A receptor that mediates the post-mating switch in *Drosophila* reproductive behaviour. Nature 451:33–37.

Zadesenets, K. S., D. B. Vizoso, A. Schlatter, I. D. Konopatskaia, E. Berezikov, L. Schärer, and N. B. Rubtsov. 2016. Evidence for karyotype polymorphism in the free-living flatworm, *Macrostomum lignano*, a model organism for evolutionary and developmental biology. PLoS One 11:e0164915.

