## Supplementary figures and images for "Effects of two seminal fluid proteins on post-mating behavior in the simultaneously hermaphroditic flatworm *Macrostomum lignano*"

### Supplementary figure 1

#### Supplementary figure 1

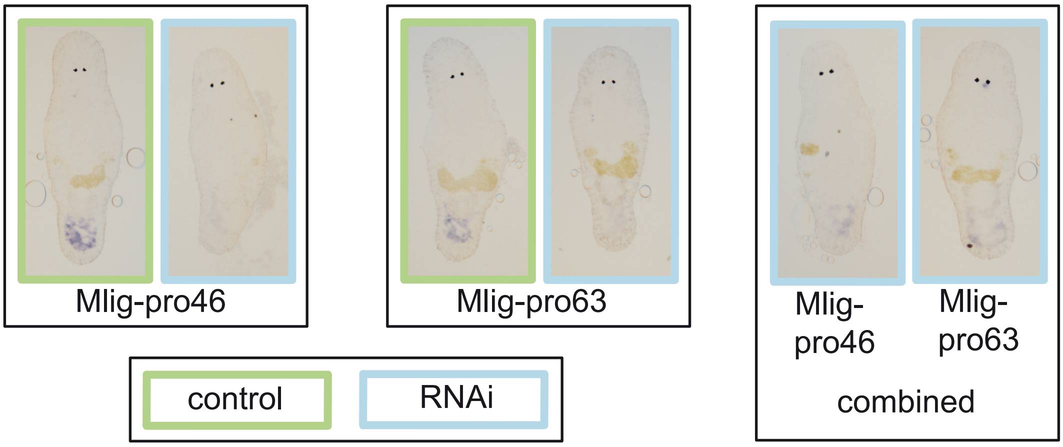
